# PoGo: Jumping from Peptides to Genomic Loci

**DOI:** 10.1101/079772

**Authors:** Christoph N. Schlaffner, Georg Pirklbauer, Andreas Bender, Jyoti S. Choudhary

## Abstract

Current tools for visualization and integration of proteomics with other omics datasets are inadequate for large-scale studies and capture only basic sequence identity information. We developed PoGo for mapping peptides identified through mass spectrometry to a reference genome to overcome these limitations. PoGo exhibited superior performance over other tools on benchmarking with large-scale human tissue and cancer phosphoproteome datasets. Additionally, extended functionality enables representation of single nucleotide variants, post-translational modifications and quantitative features.

## Main Text

Mass spectrometry (MS) and Next-generation sequencing (NGS) technologies have vastly improved our understanding of the cross-talk between genome, transcriptome and proteome, and contribute to a better understanding of the variations between healthy and diseases states. Examples are the identification of new therapeutic target kinases in breast cancer ^1^ and detection of differentially regulated pathways and functional modules potentially enabling patient stratification in ovarian cancer to inform therapeutic management. ^2^

Substantial advances in MS technologies enable more complete identification and quantification of proteomes, making these data more comparable to transcriptomics. Tools like PGx ^3^ and iPiG ^4^ to readily visualize proteomics with corresponding RNA-sequencing data on a reference genome are now increasingly indispensable. While iPiG heavily relies on the annotation format used for UCSC genes, PGx uses sample specific protein sequence databases derived from RNA-sequencing experiments and corresponding genomic coordinates. Both tools require reformatting a reference genome annotation in order to enable their mapping.

We developed PoGo to allow direct mapping to reference annotations and improve speed and quality of mapping. PoGo leverages the annotated protein coding sequences (CDS) together with a reference protein sequence database (protein-DB) to map peptides to their genomic loci. Firstly, PoGo maps the genomic coordinates of CDSs onto the protein (Figure 1), thereby connecting the protein sequences to the genomic coordinate space. Database search tools enable peptides to be identified from MS using a protein-DB. ^5^ By using the PoGo-indexed database genomic coordinates of a peptide are retrieved based on the peptide’s position within the protein (Figure 1A and Online Methods). PoGo further takes advantage of distinct attribute columns of the output file formats, such as color, to indicate uniqueness of a peptide across the genome, to show positions of post-translational modifications, to allow quantitative comparison between multiple samples and conditions linking this information to transcripts and genetic loci (Figure 2 and Online Methods). The main genome browsers, Ensembl ^6^, UCSC ^7^, and BioDalliance ^8^, however, have file size limits for direct upload insufficient for large-scale proteogenomics. Our track-hub generator application, therefore, enables seamless online visualization directly from PoGo output and is crucial for open access proteomics of large datasets.

**Figure 1.**
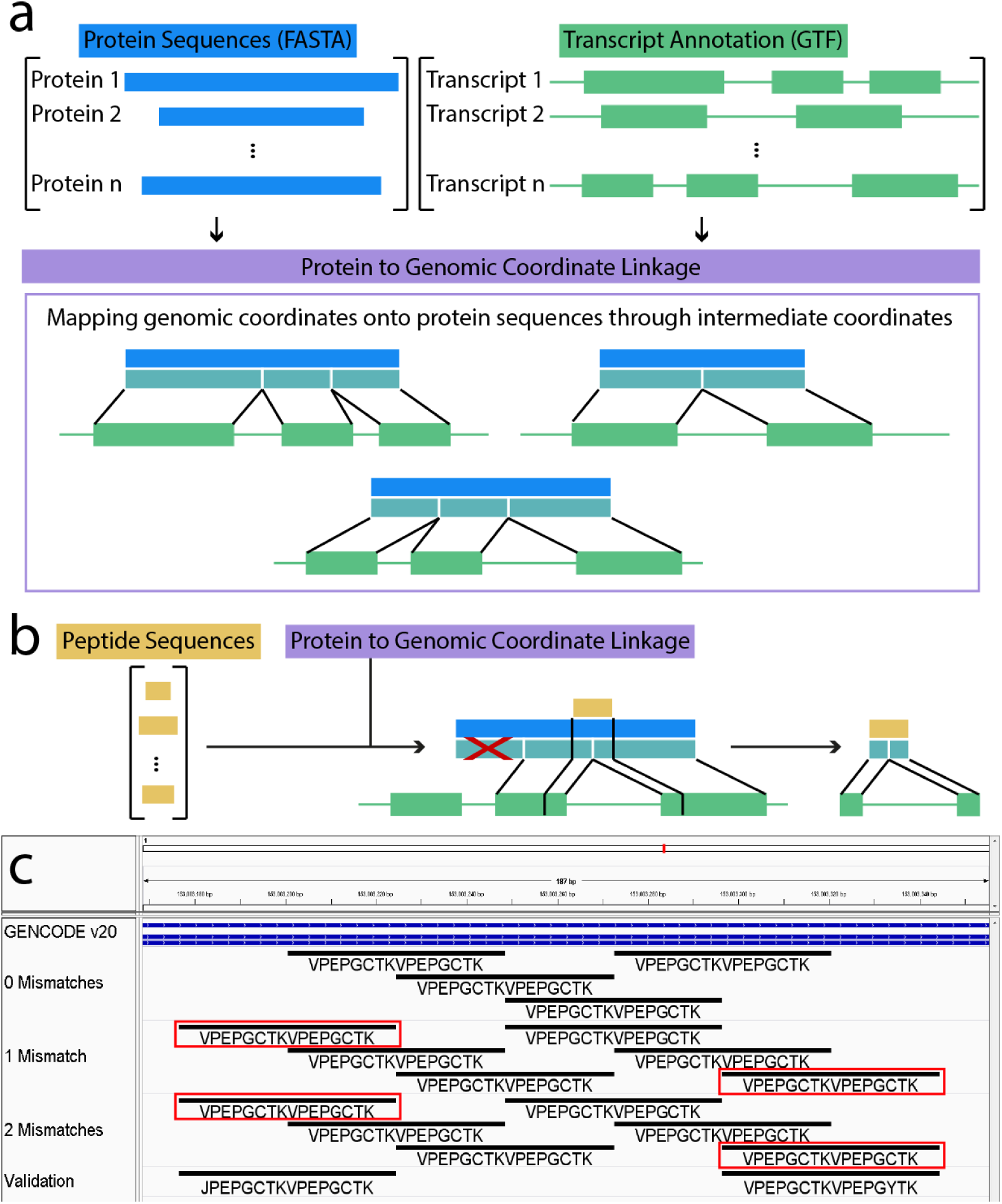
Schema of PoGo algorithm for mapping peptides through proteins to genomic loci. **(a)** Annotated protein coding transcripts in GTF format and respective translated protein sequences in FASTA format are integrated by PoGo through intermediate coordinates (turquoise), representing the exonic structure of the transcript within the protein. **(b)** Peptides, identified through searching mass spectrometry data against the protein sequence database, are mapped against the proteins. The position within the proteins then allow retrieval of overlapping coding exons and enable the calculation of the exact genomic coordinates. **(c)** Example mappings of PoGo for the overlapping repeat peptide ‘VPEPGCTKVPEPGCTK’ in a genome browser (0 Mismatches). Application of PoGo allowing for up to two mismatches results in identification of two additional repeats (1 and 2 Mismatches, red boxes) validated through mass spectrometry identified peptides of the exact sequence (Validation).

We first evaluated PoGo’s performance on large-scale datasets using the proteogenomic reanalysis of the draft human proteome maps. ^9^ We used the filtered high stringency level set comprising 233,055 unique peptides across 59 adult and fetal tissues. The mappings were derived from the gene annotation set and protein coding translation sequences for GENCODE (release 20) ^9^ as GTF and FASTA files. All tools were run with standard parameter settings and evaluated based on speed, memory usage and number of unique and correct mappings. PoGo (94 seconds) was 6.9 and 96.4 times faster than PGx (651 seconds) and iPiG (memory error after 9,064 seconds), respectively, and required 20% less memory compared to PGx (9.7 GB and 11.9 GB respectively). These data show a major improvement of speed and memory usage in addition to application with a readily available reference annotation.

In total 89% of mappings are common between PoGo and PGx. The 10.5% uniquely reported by PGx can be explained by shifting into the correct frame, indicating incorrect assignment. PoGo resulted in 89 completely unique mappings, 72 of these can be attributed to incompletely annotated transcripts (CDS start/end not found). Additionally, 17 unique mappings correspond to alternative splicing, immunoglobulin genes and multiple overlapping mappings in a repeat region. For example, the peptide ‘VPEPGCTKVPEPGCTK’ (missed cleavage between repeats of 8 amino acids) was mapped by PGx as two consecutive loci in the *SPRR3* gene (Figure S1). PoGo, on the other hand, mapped the sequence 4 times with the repeats overlapping each other (Figure 1C).

The fast and diverse mapping capabilities of PoGo, as shown above, prompted the current integration of the algorithm into the PRIDE ^10^ tool suite and soon into the OpenMS framework ^9^. This dataset also exemplifies the growing need to handle large numbers of peptides. Therefore, we have generated tissue track-hubs at two different significance thresholds from the draft human proteome maps allowing identification of genes and transcripts unique to single tissues. The scaffolding protein CASS4, for example, was only found in platelets (Figure S2). The genomic region of *RBP3*, only identified in retina, shows full peptide support for all splice junctions (Figure S3).

The large number of single nucleotide variants in individuals can affect the protein sequences and hinder identification of peptides through database searching against a reference genome. ^10^ Uniquely compared to other tools, PoGo is able to account for up to 2 non-synonymous variants in its mapping (Figure S4). Application with the draft human proteome maps allowing 1 and 2 variants resulted in a 1.5- and 60.8-fold increase in runtime (Figure S5). Unique mappings to single transcripts and single genes were reduced to 94.9% and 84.1% while the number of peptides belonging to multiple genes increased exponentially by 220.9% and 3,175.2% (Figures S5 and S6). The mapping of additional repeats of the sequence ‘VPEPGCTK’ following application with mismatches were validated through identified peptides in the sample (Figure 1C). This highlights the added value to PoGo for mapping peptides to genomic loci with potential single nucleotide variants.

To demonstrate additional PoGo functionalities we chose the phosphoproteome of high-grade serous ovarian cancer with isobaric labelling of 96 tumor samples, identifying 13,646 unique peptides with annotated phosphorylation sites (19,156 phosphopeptides). ^2^ PoGo mapped 13,617 peptides to 15,944 genomic loci in 66.9 seconds; these could not be mapped by PGx and iPiG. Only a small fraction, 0.2%, of the peptides could not be mapped due to sequence differences of the originating proteins between RefSeq and GENCODE databases. Compared to the other tools PoGo was able to use the annotated post-translational modifications and color code them (Table S1) resulting in mappings for 99.8% of the phosphopeptides with their respective localized phosphorylation sites on the reference genome (Figure S7).

PoGo also integrates peptide quantitation with genomic loci through the GCT file format. This allows comparative visualization of multiple samples in the Integrative Genomics Viewer ^11^ and enables downstream quantitative analysis. The log2-fold changes of phosphopeptides between all 69 ovarian cancer samples and the pooled reference were mapped with PoGo (Figure S7). As an example, *MAPK3* identified with multiple phosphorylated sites in a single peptide and the associated fold changes across samples are shown in Figure 2. To our knowledge PoGo is the only tool directly integrating quantitative information for peptides with genomic coordinates.

**Figure 2.**
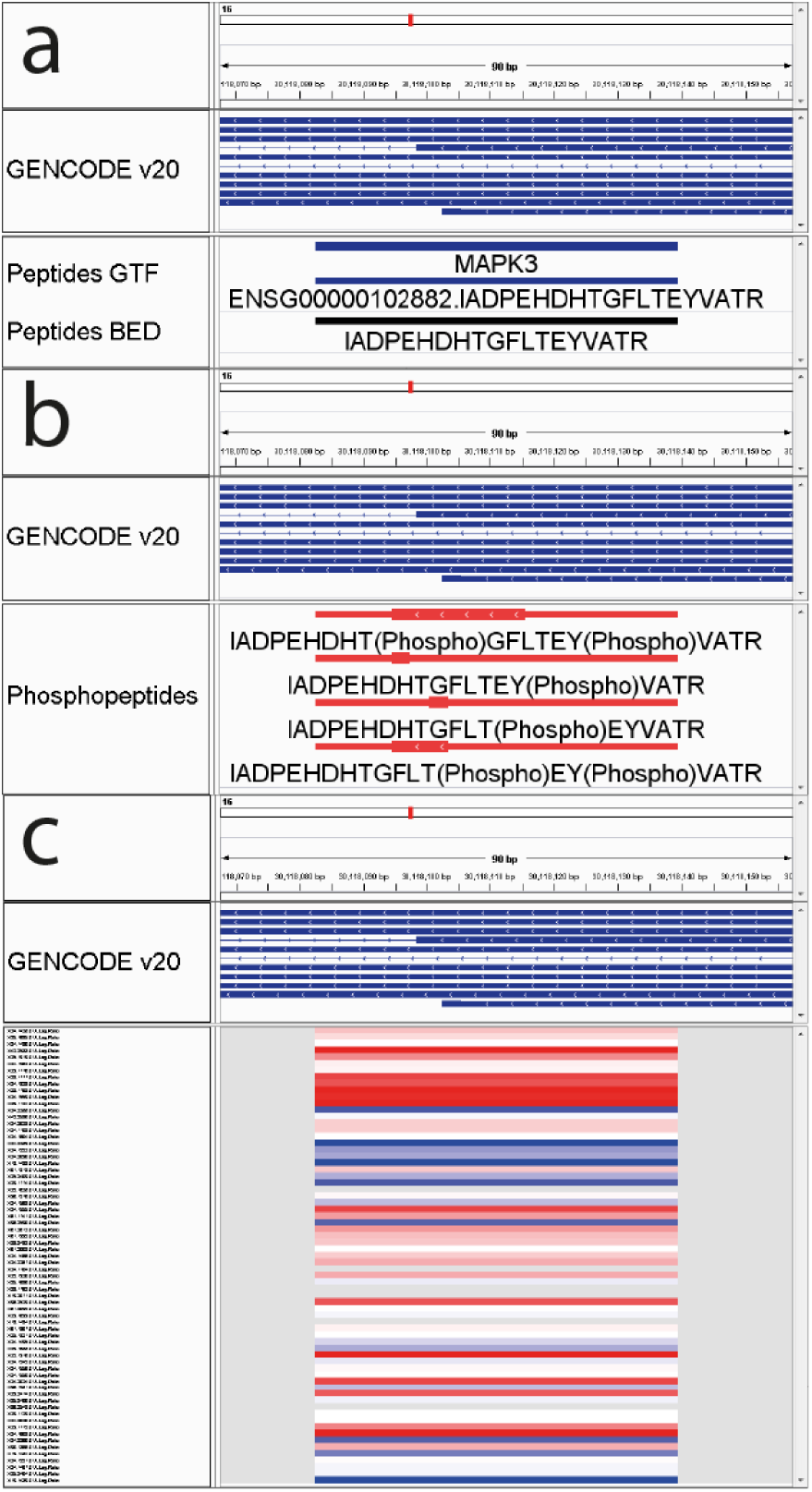
Visualization of different PoGo output formats for the peptide ‘IADPEHDHTGFLTEYVATR’ within the *MAPK3* gene. Genomic coordinates are shown at the top as x-axis. Gencode (v20) annotations of transcripts are indicated in blue. **(a)** Besides genomic location of the peptide the GTF format holds additional information, such as the gene name and gene identifier, while the BED output visualises uniqueness of the mapping across the genome. Here the black color indicates unique mapping to the gene *MAPK3*. **(b)** Genomic loci of post-translational modifications within a peptide, here phosphorylation identified through brackets in the sequence, are depicted through thick blocks spanning from the first and last modification site. The red color indicated in this output format the presence of phosphorylation. **(c)** Depiction of log2-fold changes mapped for the example peptide to the genomic location across 69 ovarian cancer samples (y-axis). High values are shown in red while blue indicates low log2-ratios.

## Discussion

Our data show that PoGo represents a major advance for peptide-to-genome mapping making it a cornerstone component of proteogenomics workflows. Although the examples used here focus on human tissue and cancer cell lines, PoGo can be applied to any proteomic study for which annotation of coding sequences in GTF format and translated sequences in FASTA format are available. The additional functionalities such as allowing up to two non-synonymous single nucleotide variants, mapping of post-translational modifications and integration of quantitation distinguish it from other tools. Semi-standardized file formats commonly used in genomics for in- and output as well as the scalability for large datasets make PoGo an indispensable component of small and large-scale multi-omics studies. The current integration into the PRIDE tool suite and our track-hub generator application promote open access proteogenomics supporting studies focusing on integration of gene, protein and post-translational modifications expression ^12^ in the future. PoGo has been developed to cope with the rapid increase of quantitative high-resolution datasets capturing proteomes and global modifications. Integration of orthogonal genomics platforms with these datasets through PoGo will be valuable for large-scale analysis such as personal variation and precision medicine studies.

## Acknowledgements

This work is funded by the National Institute of Health grant (U41HG007234) to the GENCODE project and Wellcome Trust grant (WT098051) to the Sanger Institute.

## Author Contributions

C.N.S conceived and designed the algorithms, implemented the genomic mapping algorithm, performed comparisons with other algorithms and wrote the manuscript; G.P. implemented the protein identification algorithm; A.B. and J.S.C. supervised the work and wrote the manuscript.

## Online Methods

### Software availability

PoGo is implemented in C++. Executable files for Windows and Linux, instructions for running PoGo, and explanations for each output format and their specific visual attributes are available at http://www.sanger.ac.uk/science/tools/pogo. The source code is available through https://github.com/cschlaffner/PoGo. The track hub generator application, instructions for running it, explanation of visual components of resulting track hubs, and a list of proteogenomic track hubs generated at the Wellcome Trust Sanger Institute are available at http://www.sanger.ac.uk/science/projects/proteogenomichubs. The perl source code is made available through https://github.com/cschlaffner/TrackHubGenerator.

### PoGo algorithm

PoGo is a multi-sample peptide-to-genome mapping tool taking as input tab delimited lists of peptides identified through mass spectrometry (MS) with associated number of peptide-to-spectrum matches (PSMs), quantitative value and sample identifier. PoGo also requires a reference genome annotation in the General Transfer Format (GTF) and translated protein coding sequences in FASTA format as input. The genomic coordinates of annotated coding sequences are mapped onto their respective protein sequences. Peptides identified through MS are then mapped against protein sequences accounting for up to two mismatches. The genomic coordinates for each peptide are calculated based on their position within the proteins. Each mapped peptide is additionally assigned the associated sample identifier as well as the number of PSMs and the quantitative value. Furthermore, post-translational modifications annotated in the peptide sequence are mapped to their respective genomic coordinates and color coded for the type of modification.

### Connecting protein sequences with genomic coordinates

PoGo requires protein sequences and gene annotations in FASTA and GTF format, respectively. Protein sequences have to be connected to genes and transcripts through type specific identifiers (IDs). For each protein sequence lines from the GTF file containing the transcript ID and feature-type CDS (coding sequence) are extracted. The order of exons per transcript starts with the first exon in the sequence reflecting the reading direction during translation, regardless of the strand, resulting in a reverse order of genomic coordinates for transcripts on the reverse strand. This way protein sequences and the exons match directionality. The exonic structure is mapped onto the protein sequence through construction of protein exons. Let a transcript T be a set of exons t_1_, t_2_,… t_n_ where n is the number of exons and each exon t contains the chromosome identifier, the start and end positions within the chromosome, S_t_ and E_t_ respectively, the strand on which the transcript is annotated. The corresponding protein P is defined as a set of protein exons p_1_, p_2_,… p_n_, where each protein exon p contains the start and end positions, s_p_ and e_p_ respectively, within the protein sequence so that the protein is mapped onto the transcript as f: P→T, p_i_→t_i_. For each protein in the FASTA file a map of protein exons to genomic exons is generated in PoGo.

To account for frame shifts between genomic exons t_i_ and t_i+1_ each protein exon p also holds information about the number of base pairs (bp) contributing to the codon of the first (N-term) and last (C-term) amino acid as offsets O={1,2,3}. In general, the N-term offset at the beginning of a protein defined as O(p_1_(N-term))=3 resulting in O(p_n_(C-term))=3 for complete annotations of coding transcripts. In instances where the annotation is missing a start or end codon the offsets may vary and is identified through the annotated frame. C-term offsets O(p_i_(C-term)) for each protein exon p are calculated based on the length of the genomic exon L(t_i_) and the offset of the N-term O(p_i_(N-term)) so that O(p_i_(C-term))=X=L(t_i_) mod 3-O(p_i_(N-term))+3 with the exception O(p_i_(C-term))=X mod 3 for X>3. N-term offsets of following protein exons O(p_i+1_(N-term)) are calculated so that O(p_i_(C-term))+O(p_i++1_(N-term)) mod 3=0.

### Identifying proteins of origin for input peptides

To allow fast lookup of proteins containing any given peptide PoGo creates a dictionary of words with length k (k-mer) overlapping by k-1 amino acids from the protein sequences in the FASTA input. Associated with each k-mer is a list of protein entries containing the associated protein with identifiers and the start position of the k-mer in the sequence. The dictionary is designed to consider leucine and isoleucine as equal as they are not distinguishable in MS. Peptides identified through MS are retrieved from the input file and searched against the dictionary. Thereby PoGo allows imperfect matching with up to 2 amino acid substitutions (mismatches m) to also identify proteins with potentially underlying non-synonymous single nucleotide variants. For peptides shorter than (m + 1) * k residues only the first word of length k is used and all combinations with m amino acid substitutions are generated. Each new word is looked up in the dictionary. Peptides longer than (m + 1) * k are split into consecutive k-mers and searched in the dictionary. At most m consecutive k-mers can contain amino acid substitutions leaving one word without any substitutions allowing for perfect matching in the look-up table. The presence of the peptide in each found protein then is validated taking into account the number of mismatches. The gene and transcript identifiers and the respective start position within each protein are retrieved.

### Retrieving genomic coordinates for peptides

Peptides with associated gene and transcript identifiers and the start positions within each protein are used to calculate the genomic coordinates. The length of the peptide sequence A with start position s_A_ in protein P is used to calculate the end position e_A_. To calculate the genomic coordinates for the peptide first the overlapping protein exons p are obtained so that P(A) = {x e P | s_x_ ≤ s_A_ ≤ e_x_ v s_x_ ≤ e_A_ ≤ e_x_}. Through the mapping of protein exons to genomic exons PoGo can now retrieve the genomic exons for the peptide sequence A through P(A)→T(A). The genomic coordinates then are calculated as start S_A_ = S_E_ + dS_A_ and end E_A_ = S_E_ + dE_A_ if the gene is on the forward strand or start S_A_ = S_E_ – dS_A_ and end E_A_ = S_E_ – dE_A_ if on the reverse strand with dS_A_ = (s_A_ – s_P_ – 1) * 3 + O(P(N-term)) and dE_A_ = (e_A_ – s_P_) * 3 + O(P(N-term)) – 1 denoting the distance of the genomic start and end of the peptide, respectively, from the genomic start position S_E_ of the genomic exon E.

### Mapping post-translational modifications

Besides mapping peptides, PoGo is also capable of mapping post-translational modifications (PTMs) onto the genome. Post-translational modifications are commonly annotated in the peptide sequence through round brackets containing the PSI (Proteomics Standards Initiative) name of the modification following the modified amino acid. With the position of post-translational modifications in the peptide sequence, start s_PTM_ and end e_PTM_, the mapping of the underlying peptide to the genome the above equations to calculate the genomic positions are adjusted: dS_PTM_ = (s_A_ + s_PTM_ – s_P_ - 1) * 3 + O(P(N-term)) and dE_PTM_ = (s_A_ + e_PTM_) * 3 + O(P(N-term)) – 1. Different types of PTMs are mapped separately and color coded in the output while multiple occurrences of the same PTM type, e.g. phosphorylation, within a single peptide are combined into a single mapping using the first and last PTM sites.

### Adding quantitative information for multiple samples

To allow visualization of quantitative information for peptides on a genome, PoGo records this type of information. Peptide and sample pairings may only occur once in the input file uniquely identifying a quantitation value. PoGo stores the tuples of sample identifier, quantitative value and the number of peptide to spectrum matches (PSMs) for each peptide. This information is used in the different output formats to allow comparative analysis.

### Generating different output formats

PoGo generates output in three formats commonly used in genomics. The first and central output format of PoGo is BED. This format stores each mapped peptide as a single line of twelve tab delimited columns. Besides chromosome coordinates, the peptide sequence, strand as well as start and end coordinates of a thick block the start positions and lengths of peptide blocks mapping to genomic exons are included. Additionally, BED files support individual coloring of each feature. PoGo utilizes this in two different forms. Firstly, in the general peptide centric output of PoGo peptides are colored based on their uniqueness within the genome. Peptides unique to a single transcript are colored in red while peptides shared between multiple transcripts of a single gene are shown in black. Peptides mapping to multiple genes are indicated by their grey color. Secondly, PoGo also generates a separate BED file for peptide forms with post-translational modifications. In this instance the thick block element is used to indicate the position of the post-translational modification. Two or more modifications of the same type within a single peptide sequence are collapsed to indicate the range between the first and last modification site. The coloring of the uniqueness per peptide in the genome is substituted to accommodate color coding of post-translational modifications.

The second file format supported by PoGo for mapped peptides is the general transfer format (GTF). PoGo redefines some of the feature types to accommodate mapping of peptides. The feature type ‘transcript’ is used to indicate a mapped peptide while the feature type ‘exon’ indicates the concrete mapping of the peptide to underlying genomic exons. PoGo additionally stores information such as the gene identifier, name and biotype for the gene as well as the number of peptide-to-spectrum matches (PSMs) and quantitative values for each sample in which the peptide was identified.

For comparative or quantitative analysis PoGo generates the output format GCT which can be visualized in the Integrative Genomics Viewer (IGV). ^11^ This third format is similar to a matrix with rows identifying a peptide with genomic mapping and columns identifying a sample. Each cell holds the quantitative values associated with the peptide and the sample given in the input file.

### Human tissue data

High-resolution MS data from 59 fetal and adult human tissues were used for the validation of PoGo. The raw data of these draft human proteome maps were generated by the Pandey lab ^13^, the Kuster lab ^14^, and Cutler lab. ^15^ All three datasets were combined and reprocessed by Wright et al. Wright, Mudge, Weisser, Barzine, Gonzalez, Brazma, Choudhary and Harrow ^9^ The data were retrieved in a tab delimited format combining all results from mzid files available from PRIDE Archive. ^10^ Identifications were filtered to the highest stringency level described in Wright et al. ^9^ for identification of novel coding regions (q-value ≤ 0.01 (1% FDR), a PEP of ≤ 0.01, peptide length between 7 and 29 residues, full tryptic peptides, a maximum of two missed cleavages).

### Phosphoproteomic ovarian cancer data

We applied PoGo to isobaric labelled phosphoproteome data from an ovarian tumor study comprising 69 samples. ^2^ Phosphopeptides with associated iTRAQ quantitation were downloaded as tab separated file from https://cptac-data-portal.gorgetown.edu. Lower case characters (s, t and y) in the peptide sequence showing phosphorylation were substituted by upper case characters followed by the PSI name of phosphorylation in brackets.

### Protein sequences and gene annotation and PoGo settings

The annotation of human genes in GTF format and the corresponding protein coding sequence translation as FASTA files were downloaded for GENCODE v20 ^9^ from http://www.gencodegenes.org. Gene and transcript identifiers were set as “ENSG” and “ENST” for genes and transcripts, respectively, followed by 11 digits and the word length for k-mers was set to 5 amino acids. For post-translational modifications 10 biologically relevant types were chosen for easy discriminability of the color code (Table S1).

### Comparison of algorithms for performance evaluation

For the human tissue and the ovarian cancer phosphoproteome data PoGo’s performance was compared against PGx ^3^ (downloaded from https://github.com/FenyoLab/PGx) and iPiG ^4^ (downloaded from https://sourceforge.net/projects/ipig/), two standalone tools available to map peptides to their corresponding genomic coordinates. Each dataset was formatted using in-house scripts in R and perl to fit the required input format for each tool. Each program was run using default parameters and the minimum number of required input files. To compare the mappings between the tools, we marked as equal when chromosome name, start and end positions, the exon starts and lengths as well as the peptide sequence were the same. Frameshifts then were identified amongst unique mappings per tool through shifting either start or end position by up to two base pairs and comparing those to the consensus mappings. Remaining unique mappings of the tools then were examined manually by comparing the peptide sequence to the translated sequence of the respective genomic coordinates in the IGV browser. ^11^

### Generating track hubs

Track hubs were generated to visualize different aspects of the human proteome maps. The data was filtered to two stringency levels resulting in two sets. The first result set was filtered to a standard significance (q-value of ≤ 0.01 (1% FDR), a PEP of ≤0.05 and a minimum peptide length of 7 residues) while the highest stringency level mentioned in Wright, Mudge, Weisser, Barzine, Gonzalez, Brazma, Choudhary and Harrow ^9^ (q-value ≤ 0.01 (1% FDR), a PEP of ≤ 0.01, peptide length between 7 and 29 residues, full tryptic peptides, a maximum of two missed cleavages) was applied to the second set. Additionally, each set was split into subsets for individual tissues, resulting in 60 files per set. PoGo was run with default parameters using the property of passing a comma separated list of input files to be mapped separately. The Track-Hub Generator application then was run using the 60 output files in BED format to generate two track hubs; one for each significance level filter. Folders and files required for track hubs are generated automatically. The script ‘fetchChromSizes.sh’ and tool ‘bedToBigBed’ from UCSC (both downloaded from http://hgdownload.cse.ucsc.edu) ^16^ are used in the Track-Hub Generator to create binary files from the original BED files used for track hubs. The generated track hubs are accessible through ftp and http via http://www.sanger.ac.uk/science/projects/proteogenomichubs.

